# Host specificity of fungal functional groups covaries with elevation: implications for intraspecific plant interactions

**DOI:** 10.1101/2025.05.24.655939

**Authors:** Abigail S. Neat, Felipe Albornoz, Kyle A. Gervers, Posy E. Busby, F. Andrew Jones

## Abstract

Plants of the same species can harbor shared enemies, resulting in indirect, intraspecific competition. When this competition is strong, there are fitness costs associated with growing near a same-species individual, termed Conspecific Negative Density Dependence (CNDD). Host-specific microbial pathogens are known mediators of this type of indirect, intraspecific plant competition, yet how these plant-microbe interactions vary with environmental context is less explored. Further, microbial pathogens and mutualists may jointly contribute to the relative strength of intraspecific competition, yet most studies focus only on pathogens. We use ITS metabarcoding to characterize soil and foliar fungal functional groups associated with individual trees of three dominant conifers in forests where the strength of CNDD is known to be greater at lower elevations relative to higher elevations. We hypothesize that plants found in the mesic low elevation forests accumulate more host-specific fungal pathogens compared to plants found in the more xeric high elevation forests. Conversely, we hypothesize that plants found in the high elevation forests accumulate more host-specific mutualists compared to those found at low elevations. We test these hypotheses by evaluating three metrics of plant-associated fungal functional groups – alpha diversity, relative abundance, and host specificity – that together address the degree to which a plant recruits particular groups. We find the diversity, relative abundance and host specificity of soil fungal pathogens to decrease with elevation. In contrast, we find ectomycorrhizal fungal host specificity, but not diversity or relative abundance, to increase with elevation. Aboveground, we find a key foliar pathogen *Nothophaeocryptocus gaeumanii* to support our hypotheses as a dominant, and highly host specific pathogen that is enriched at low elevation sites. Our results indicate that across a climate gradient known to negatively covary with CNDD strength, soil pathogens recruitment is most prevalent in the mesic climates and, to a lesser extent, ectomycorrhizal recruitment is most prevalent in the xeric climates. Together these findings suggest that both pathogenic and mutualistic fungal symbionts could contribute to landscape level variation in competitive, intraspecific plant interactions.

## Introduction

Plant-microbe interactions range from mutualistic to antagonistic (Mendes, Garbeva, and Raaijmakers 2013; Schirawski and Perlin 2018) and can strongly affect plant populations, communities, ecosystem structure, diversity, and dynamics (Bever, Platt, and Morton 2012; Gilbert and Parker 2023; Reynolds et al. 2003). Pathogenic microbes can colonize all plant tissues including roots, leaves, and stems (Beattie and Lindow 1995; Castello, Leopold, and Smallidge 1995). Host-specific pathogenic microbes are hypothesized to facilitate indirect, apparent competition between plants of the same species by accumulating in areas with high densities of conspecific individuals and beneath the canopies of adult individuals (Janzen 1970; Connell 1971; Wright 2002). When host-specific enemies act in a density dependent manner, conspecific negative density dependence (CNDD) can be a stabilizing mechanism that regulates population growth and maintains species diversity by slowing the rate of competitive exclusion in communities (Bagchi et al. 2014; Forrister et al. 2019; LaManna et al. 2024; Liu et al. 2022; Mangan et al. 2010).

Plants also host mutualistic microbes that may destabilize community dynamics and reduce plant species diversity (Bever, Platt, and Morton 2012; McGuire 2007; Zahra, Novotny, and Fayle 2021). Almost all plant species associate with mycorrhizal fungi, which can enhance plant water uptake, nutrient acquisition (Chalot and Brun 1998; Smith and Read 2008), and the ability to overcome biotic and abiotic stresses (Pozo and Azcón-Aguilar 2007). Aboveground, foliar microbes have been shown to both directly and indirectly confer tolerance to biotic and abiotic stressors, such as drought, salinity, and pathogens (Busby, Ridout, and Newcombe 2016; Rodriguez et al. 2008). Microbial mutualists can increase an adult plant’s fitness and the chances of conspecific offspring survival to reproductive age (Arnold et al. 2003; Koide and Dickie 2002; Teste and Simard 2008), resulting in the local recruitment of conspecific seedlings and weakening CNDD (Bever, Platt, and Morton 2012; McGuire 2007). Therefore, both plant mutualists and enemies may differentially contribute to CNDD strength.

Plant-microbial interactions have been demonstrated to be context dependent and vary based on environmental conditions (Dudenhöffer, Luecke, and Crawford 2022; David et al. 2020; Hoeksema et al. 2010). Moist, mesic climates are documented to harbor pathogens and facilitate plant-pathogen interactions (Spear, Coley, and Kursar 2015; Connell 1971; Milici et al. 2020; Rutten and Gómez-Aparicio 2018). In contrast, mutualistic plant-microbial interactions have been found to be especially important in dry, arid, and drought-prone climates (Bingham and Simard 2012; Rutten and Gómez-Aparicio 2018; David et al. 2020), and are hypothesized as a mechanism for plant and their associated microbial communities to persist in climates challenged with abiotic stress (Bertness and Callaway 1994; Rodriguez et al. 2008; David et al. 2020). In parallel, past research has also demonstrated the environmental context dependency of intraspecific plant interactions, with greater competition observed in wet, mesic climates as compared to dry, arid climates both at local (García-Cervigón et al. 2013; LaManna et al. 2022) and global scales (LaManna et al. 2017). However, connections between intraspecific competition (ie CNDD), the microbes potentially mediating these interactions, and climate, remain sparse.

Fungi include significant plant pathogens and mutualists, and are therefore an important subset of the plant microbiome regulating plant species distributions (Bagchi et al. 2014; Bell, Freckleton, and Lewis 2006; McGuire 2007). Until recently, studies investigating the contributions of fungi to CNDD strength have focused primarily on soil pathogens. However, plants ubiquitously associate with diverse microbes beyond soil pathogens including mycorrhizal fungi and foliar fungi. Both of these subsets of the plant microbiome have been shown to generate plant feedback (Teste et al. 2017; Whitaker et al. 2017) yet their relative contributions to CNDD strength remain less explored (see Chen et al. 2019). Further, experiments investigating the role of fungi in shaping plant species distributions have been executed primarily in the tropics or at single sites within forest stands (Bagchi et al. 2014; Chen et al. 2019; Mangan et al. 2010). As a result, gaps remain in our understanding of (1) how functional groups of plant-associated fungal communities differ across environmental gradients, (2) how this variation may in turn result in landscape level differences in CNDD strength and (3) the extent to which mutualists and foliar fungal communities might contribute to these patterns in plant species distributions.

In this paper, we integrate theory on the environmental context dependency of both intraspecific plant competition and plant-microbial interactions to explore the contributions of microbes to patterns of CNDD. LaManna et al. (2022) characterized the strength of CNDD for the growth and survival of 18 tree species across a 1,000 meter elevation and climate gradient in montane coniferous forests in Oregon, USA. For all 18 species examined, CNDD was more negative for tree individuals within mesic and warmer lower elevation sites relative to the arid and colder higher elevation sites. Attenuating CNDD at higher elevations could result from (1) reduced pathogen accumulation in these high elevation forests compared to the lower elevation forests, or (2) reduced pathogen accumulation and greater mutualist accumulation jointly (Figure 1). Here, we use ITS metabarcoding to examine host and environmental drivers of foliar and soil fungal communities associated with individual trees of three dominant conifers found across the same gradient. We evaluate three metrics of the host-associated functional groups (hereafter ‘functional guilds’) which together represent the degree to which a plant recruits the particular guild: alpha diversity, relative abundance, and host specificity. We hypothesize that plants found at low elevations will exhibit more host-specific pathogen accumulation relative to those found at high elevations; in contrast, plants found at high elevations will accumulate more host-specific mutualists relative to those found at low elevations.

**Figure 1.**
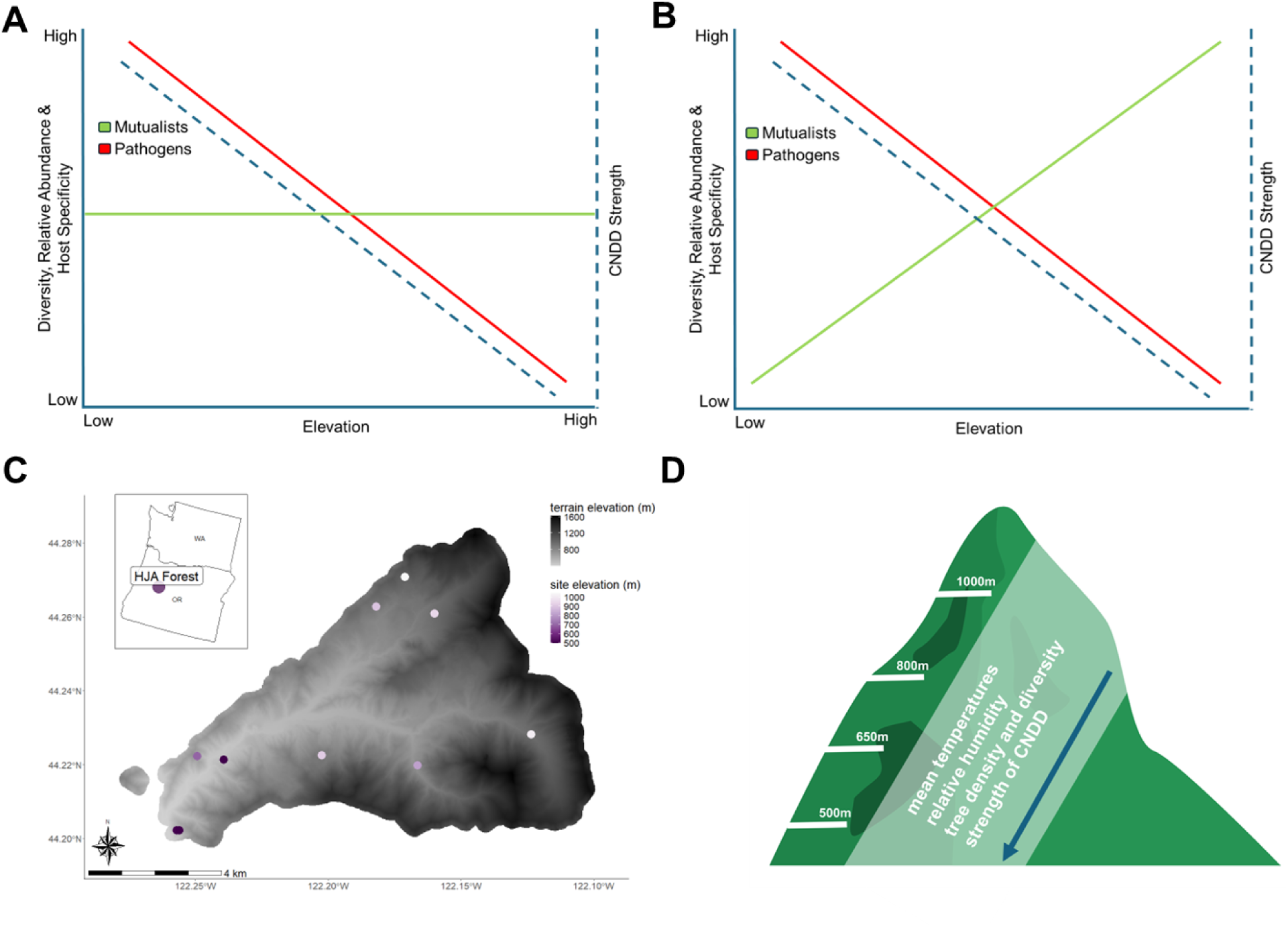
(A) and (B) outline how fungal symbiont communities may contribute to the observed negative relationship between CNDD and climate. Map (C) and conceptual figure (D) of the climate gradient present at the HJ Andrews Experimental Forest. Two potential expectations for diversity, relative abundance and host specificity of fungal guilds across the gradient, with either outcome supporting the trends CNDD strength observed by LaManna et al (2022) (A,B). Map displaying the 10 study sites, with the elevation of each site indicated by the purple color gradient (C). Conceptual figure showing attributes of the climate and forest that are directly related to elevation and are greater at lower elevation sites (D).

## Materials and Methods

### Study Sites

The study was conducted across 10 forest stands located within HJ Andrews (HJA) Experimental Forest in the Cascades Mountain Range of Western Oregon (44.21°N, –122.25°W) (Figure 1). These sites are ¼ hectare reference stands that were established in 1971 to study plant communities across elevation and over time, and vary from 500 to 1050 m in elevation (Hawk and Biome 1978). The elevation gradient also represents a climate gradient, with sites decreasing in average annual temperature (range = 8.6℃ to 10℃), relative humidity (range = 57% to 63%), and mean annual precipitation (range = 2066 mm to 2139 mm) (Figure 1). Tree species diversity is greater at the lower elevation, mesic, and warmer sites relative to the higher elevation, drier, and colder sites (LaManna et al. 2022).

### Focal Species and Sample Collection

We characterized foliar and soil fungal communities associated with individual trees representing three focal study species, Douglas-fir (*Pseudotsuga menziesii* var *menziesii*) (PSME), Western Hemlock (*Tsuga heterophylla*) (TSHE), and Pacific Yew (*Taxus brevifolia*) (TABR). These three species are among the most dominant in Pacific Northwest forests of North America, and their ranges span the elevation gradient. These species also have distinct life history strategies and mycorrhizal fungal associates. *P. menziesii* is the most abundant conifer across the gradient and an early successional species (McArdle and Meyer 1961). *T. heterophylla* is a secondary colonizer, exhibits slower growth relative to *P. menziesii*, and is shade tolerant (Burns and Honkala 1990). *T. brevifolia* is a late successional shade tolerant species, often rare and extremely slow growing relative to the other two species (Bolsinger and Jaramillo 1995). *P. menziesii* and *T. heterophylla* form ectomycorrhizal fungal associations whereas *T. brevifolia* forms both ectomycorrhizal and arbuscular mycorrhizal associations (Griffiths et al. 1995; Schoenberger and Perry 1982).

We collected paired foliar and soil samples from individual *P. menziesii*, *T. heterophylla* and *T. brevifolia* tree hosts located across the 10 sites. Within each reference stand, we haphazardly collected samples from trees associated with the three species in our study (average number of trees sampled per stand = 6, range = 4-12). Soils were collected by taking four soil cores of the top 20 cm of soil in each cardinal direction around the dripline of the focal tree. Soil samples from each tree were pooled, homogenized, and stored on dry ice for 24 hours during transport. The soil corer and collection equipment were surface sterilized with bleach and ethanol between samples, focal trees, and sites. After 24 hours, soil samples were stored in a –80°C freezer.

We collected foliar samples on two consecutive days in May 2017 using a research slingshot to obtain live needles from tree canopies, and ensuring the needles were collected before reaching the ground. Needles from an individual tree were combined in a 15 ml tube, stored in the field on dry ice then transferred to a –80°C freezer. Before DNA extraction, needles were lyophilized, and ground to a powder using a bead mill.

### Nutrient Analyses

Soil and foliar nutrient analyses were performed in the Soil Health Lab at Oregon State University, Corvallis OR. Carbon (C) and nitrogen (N) were measured using dry combustion on Vario Macro Cube (Elementar, Langenselbold, Germany), and organic matter (OM) was calculated using total organic carbon * 2 (Pribyl 2010). pH was measured in a 1:1 soil:water ratio on a HI5522 benchtop meter (Hanna, Woonsocket, Rhode Island). Calcium (Ca), magnesium (Mg), and potassium (K) were quantified with a 1M ammonium acetate extraction measured on 5110 ICP-OES (Agilent, Santa Clara, California). Boron (B), copper (Cu), iron (Fe), manganese (Mn) and zinc (Zn) were quantified using DTPA-sorbitol extraction measured on 5110 ICP-OES (Agilent, Santa Clara, California).

### Climate Data

The temperature variables used in this study include mean summer (July to September) temperatures, maximum summer temperatures, minimum temperature of the coldest month, mean spring temperatures (April to June), maximum summer temperatures, and minimum summer temperatures. All data were collected between 2009 and 2018 using understory temperature loggers (1.5m tall) at each of the study sites (LaManna et al. 2022). These variables were chosen because they have been previously demonstrated to determine the strength of CDD across the gradient (LaManna et al. 2022).

### DNA Extraction, Amplification, and Sequencing

For soil samples, we extracted DNA from ∼ 250 mg of soil using the PowerSoil DNA isolation kit (#12888-100, QIAGEN) following manufacturer’s protocol. For foliar samples, we extracted DNA from ∼ 20 mg of freeze-dried material using the Plant DNeasy kit (#69204, QIAGEN) following manufacturer’s protocol.

We amplified the ITS1 region using ITS1F (Gardes and Bruns 1993) and ITS2 (White et al. 1990) primers. The PCR reaction for both the soil and foliar samples was as follows: 1X PCR buffer (with MgCl_2_), 1.5 mM MgCl_2_, 0.2 mM dNTP mix, 0.2 uM of each primer (ITS1F and ITS2), and 2 U/rxn of Platinum Taq (#10966018, ThermoFisher Scientific). Two microliters (μl) (l) of genomic DNA were added to each reaction for a total volume of 25ul. Polymerase Chain Reaction (PCR) amplification was conducted on a T100 Thermocycler (Bio-Rad, USA) with the following settings: 95°C for 5 minutes, and 34 cycles of 95°C for 30 s, 72°C for 2 min, and finally 72°C for 5 min. Amplicon clean-up, ligation of Illumina NEXtera XT indices, and final library purification were conducted by the Center for Quantitative Life Sciences (CQLS) at Oregon State University. We sequenced the amplicons using the Illumina MiSeq platform with 2×300-bp paired-end reads.

### Bioinformatics

We demultiplexed reads and removed the primers using Cutadapt (Martin 2011). We then filtered and trimmed, denoised, dereplicated, merged read pairs, and removed chimeras from reads using ‘dada2’ (Callahan et al. 2016). We extracted the ITS1 region using ITSxpress (Rivers et al. 2018) to produce a tree host site-by-species Amplicon Sequence Variant (ASV) table. We then clustered ASVs into Operational Taxonomic Units (OTUs) using the R package ‘DECIPHER’ in which reads that share at least 95% sequence similarity are grouped into a single OTU (Wright 2016). Prior to taxonomy assignments, we used the *2023-07-18 UNITE all eukaryote* reference database to remove reads from the dataset that were not in kingdom fungi or were unknown at the kingdom level (Abarenkov et al. 2024). We then assigned taxonomy to the remaining reads using the *2024-03-25 fungal UNITE+INSD* reference database, which were then used in downstream analyses (Abarenkov et al. 2024). Both steps of the taxonomy assignments were performed using the *assigntaxonomy* function in ‘dada2’ (Callahan et al., 2016).

### Statistical Analyses

We split the soil and foliar fungal community datasets to be analyzed and interpreted separately. Unless otherwise stated, analyses were performed on both datasets. The soil dataset included 66 individual tree hosts; the needle dataset, 65. We performed rarefaction to 25,303 reads, to standardize the sampling depth to the sample with the least number of reads between the two datasets (Schloss 2023) (Appendix S3: Figure S1). This was executed using the *rrarefy* function from the ‘vegan’ R package (Oksanen 2010) to generate a new community matrix that had been subsampled to the standardized sampling depth. The process was then repeated iteratively 1,000 times, and the matrices averaged to create a final community matrix to be used for subsequent analyses.

The species matrix (soil = 66 trees × 1,847 OTUs, foliar = 65 trees x 1,072 OTUs) contains counts of the number of rarefied sequence reads for each OTU per focal tree. The environmental matrix (soil = 66 trees × 44 variables, foliar = 65 trees x 44 variables) contains information on each tree host (species, site, elevation), measurements of the foliar nutrient composition (C, N, P, etc.) and soil properties (moisture, organic matter, nutrients, etc) associated with each sample. We were unable to obtain nutrient information for some samples because insufficient material remained after DNA extraction. In cases in which either foliar or soil nutrients were unknown, we used the median value associated with each tree host species. For each nutrient, an average of 12 samples (12/131 or ∼ 9%) were missing and required this median value assignment.

The guild matrix (soil = 1,847 OTUs x 4 guilds, foliar = 1,072 OTUs x 2 guilds) contains information on OTU fungal guild classification obtained using a combination of the R packages ‘FungalTraits’ (Põlme et al. 2020) and ‘FUNGuild’(Nguyen et al. 2016). In brief, taxa were first sorted into guilds using the FungalTraits database. Those taxa that were not assigned a guild were then passed through the FUNGuild database, and ‘probable’ and ‘highly probable’ assignments were used to classify the remaining taxa not yet assigned a guild. We used both databases, because there are assignments that are present in FungalTraits and not FUNGuild, and vice versa. This final matrix contains binary data indicating an OTU’s guild affiliation, where ‘1’ signifies known guild affiliation and ‘0’ signifies non-affiliation or unknown affiliation status.

### Community Analysis

We calculated fungal community richness and the Shannon diversity associated with each focal tree host using the R package ‘vegan’ (Oksanen 2010). We also used ‘vegan’ and its PERMANOVA (Anderson 2001) function *adonis2* (permutations = 999, by = ‘margin’) to test the roles of climate, tree host species, soil properties, foliar nutrients, and soil pH in shaping fungal community composition. The data were visualized with the ‘ggplot2’ package (Wickham 2009). To reduce dimensionality of the climate, soil properties, and foliar nutrients data, we first scaled all variables, then performed three separate Principal Component Analyses (PCAs). The following variables were included in the climate PCA: mean summer temperatures, maximum summer temperatures, minimum temperature of the coldest month, mean spring temperatures, maximum summer temperatures, minimum summer temperatures. Maximum spring temperatures and maximum summer temperatures both negatively correlated with the first principal component. The first principal component (hereafter, climate) explained 82.3% of total variation (Appendix S3: Figure S2) with lower values corresponding to lower elevation sites and higher values corresponding to higher elevation sites (Appendix S3: Figure S2). The soil properties PCA included soil moisture, organic matter content, C:N ratio, cation exchange capacity, and soil nutrients. The first principal component correlated with soil manganese and iron and explained 53.2% of the total variation (Appendix S3: Figure S2). The foliar chemistry PCA included foliar nutrients (C, N, Ca, Mn, etc) and foliar manganese correlated with the first principal component which explained 91.1% of total variation (Appendix S3: Figure S2). In all three cases, only the first component of the PCA was used in the PERMANOVA analyses. The soil community models did not include the foliar nutrients variable, and the foliar community models did not include the soil properties variable. PERMANOVA model comparison was evaluated using the Akaike information criterion corrected (AICc) forward model selection (Appendix S2: Tables S1). For cases in which the lowest AICc values were within 2 units of each other, we selected the model with the lowest AICc values.

To examine and visualize the relationship of fungal community composition and the environmental variables, we used non-metric multidimensional scaling (NMDS) with Bray-Curtis distance measures. We executed the ordination with the R package ‘vegan’ and its function *metaMDS* (Oksanen 2010), and evaluated the data in two dimensions.

### Fungal Guild Analysis

The fungal guilds included in this analysis were plant pathogens, arbuscular mycorrhizal fungi (AMF), and ectomycorrhizal fungi (EMF). Foliar mutualists were excluded from the guild level analyses because this group is only well defined for endophytic fungi in the family Clavicipitaceae and their associations with grasses (Clay 1988), and are otherwise comparatively understudied (Sieber 2007). For these foliar fungal communities, we instead supplemented the analyses described below with simple linear regression analyses of how the relative abundance specific taxa are associated with climate. Otherwise, all analyses were performed at the guild level. We used three separate metrics to evaluate host recruitment of each guild: alpha diversity, community weighted mean (CWM) (Legendre, Galzin, and Harmelin-Vivien 1997), and host specificity. For the diversity analysis, we calculated a site level Shannon’s diversity index for each guild associated with each focal tree. For the CWM analyses, each site received a CWM value that represents the relative presence of each fungal guild across the focal tree hosts. For example, a community with a greater proportion of pathogens would have a higher pathogen CWM value. This value was determined using the trait matrix (described above) and the *makecwm* function in the R Package ‘ecole’. Finally, we created a host specificity index for each fungal guild associated with each focal tree. Here we define host specificity as the degree to which a microbial species is associated with a certain host, or rather, is more widely associated with many host species (Christian et al. 2017). In brief, we generated the index by first assigning each taxon a specificity value derived from the permutation test of the indicator species analysis (Dufrêne and Legendre 1997). These values range from 0-1, with values closer to 0 indicating a generalist taxon and values closer to 1 indicating a host specific taxon. We then used the specificity values associated with each taxon to perform a CWM analysis. This analysis generated focal host level specificity indices representing the degree of specificity for each guild community as a whole. More information on calculating the specificity index can be found in Appendix S4. For each fungal guild, we used multiple linear regressions and AICc forward model selection (Appendix S2: Tables S2-S7) to test the relationship between the environmental variables and our three metrics (alpha diversity, CWM values, host specificity). Consistent with the above PERMANOVAs, the soil community models did not include the foliar nutrients variable, and the foliar community models did not include the soil properties variable. For cases in which model scores were within two units of the lowest AICc score, these models were averaged for parameter estimates (Symonds and Moussalli 2011).

## Results

### Fungal community and guild generation

After trimming primer sequences and removing control samples, we obtained 15,325,090 reads from 131 total samples. Through filtering, alignment, chimera removal, denoising, and extracting the ITS1 region, reads were filtered down to 11,773,360 reads of 26,430 ASVs. The processes of clustering to 95% similarity generated 8,859 OTUs. We then removed reads that were unknown to kingdom level, not in the kingdom Fungi, or had fewer than 20 reads in the data set, which together resulted in 2,586 OTUs from 131 samples for downstream analyses.

We found 1,847 OTUs across 66 tree host sample units in the soil fungal community dataset. Of these OTUs, 665 were classified as saprobes, 74 as pathogens, 464 as EMF and 11 as AMF. We found 1,072 OTUs across 65 sites in the foliar fungal community dataset. Of these OTUs, 374 were classified as saprobes and 122 as pathogens. To address our hypotheses, we focused the analyses on pathogen and mutualists (EMF and AMF) guilds. Analyses associated with saprobes can be found in the appendices.

### Community composition, richness and diversity

Soil fungal community richness was negatively associated with climate PCA axis 1 (β = –9.35, p = 0.002) (Figure 2, Appendix S1:Table S1). This climate variable is correlated with elevation, indicating that soil fungal richness was higher at lower elevation sites compared to high elevation sites. We found no significant association between soil community diversity and the climate PCA axis 1 (Figure 2, Appendix S1:Table S1). In our analysis of soil fungal community composition, climate was the only factor to significantly impact community composition (PERMANOVA, R^2^ = 0.08, p = 0.001, F = 5.34) (Figure 2, Appendix S1: Table S2).

**Figure 2.**
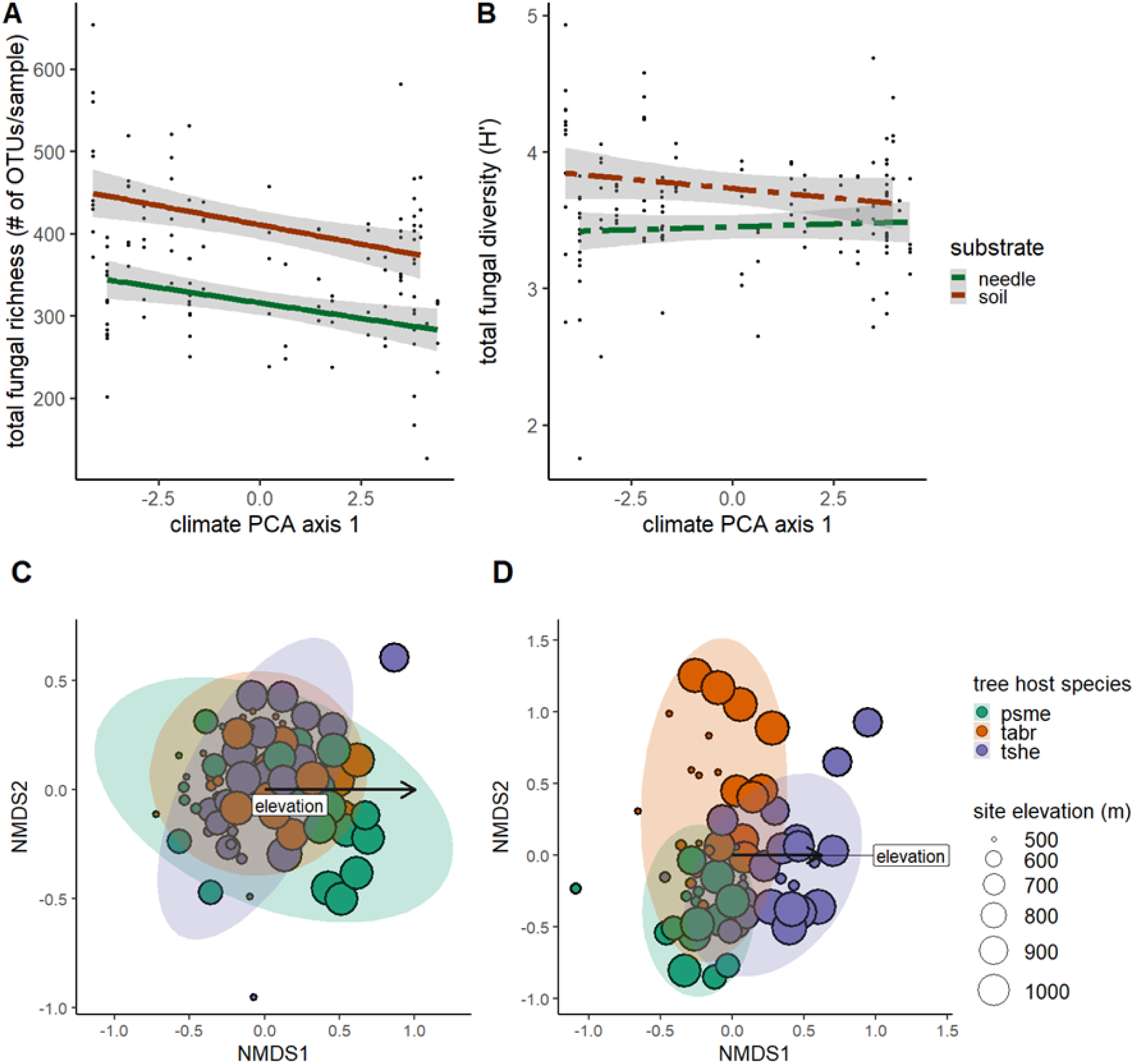
Total fungal richness (A) and the Shannon Diversity Index (B) show both soil and needle fungal communities have greater richness associated with smaller climate values and thus lower elevation sites. NMDS ordinations show that (C) soil fungal communities are structured by climate gradient (R^2^ = 0.08, p = 0.001) while (D) foliar fungal communities are structured primarily by tree host species (R^2^ = 0.17, p = 0.001) and secondarily by the climate gradient (R^2^ = 0.07, p = 0.001). For (A) and (B), the x-axis represents the ‘climate’ variable derived from the first principal component of a PCA. The ‘climate’ variable is directly related to elevation with negative values associated with lower elevation sites and positive values associated with higher elevation sites. Significant relationships have solid lines (p < 0.05), while insignificant relationships have dashed lines. For both ordinations, sample units are in species space using Bray-Curtis distance measure (C,D). The overlaid vector represents correlation of environmental variables elevation to the ordination. The ellipses indicate a 95% confidence interval for the multivariate probability distribution.

We found similar patterns for foliar fungal richness and diversity. Richness, but not diversity, was negatively associated with climate PCA axis 1 (β = –7.49, p = 0.003) (Figure 2, Appendix S2: Table S2). Thus foliar fungal community richness declined across the elevation gradient. Unlike patterns in soil fungal community composition, tree host species most strongly influenced foliar community composition (PERMANOVA, R^2^ = 0.17, p = 0.001, F = 6.83), followed by climate (PERMANOVA, R^2^ = 0.07, p = 0.001, F = 5.47) (Figure 2: Appendix S1: Table S2).

### Fungal guilds and the climate gradient

For soil fungal pathogens, all three of our metrics for assessing diversity, relative abundance, and host specificity were correlated with climate PCA axis 1, and all tests indicated a decline across the elevation gradient. Soil pathogen alpha diversity (β = –0.07, p = 0.001), CWM values (β = –0.002, p = 0.002), and host specificity (β = –0.008, p = 0.001) were all negatively associated with the climate variable (Figure 3, Appendix S1: Table S3). These findings support our hypothesis that pathogens are more enriched at lower elevations relative to higher elevations.

**Figure 3.**
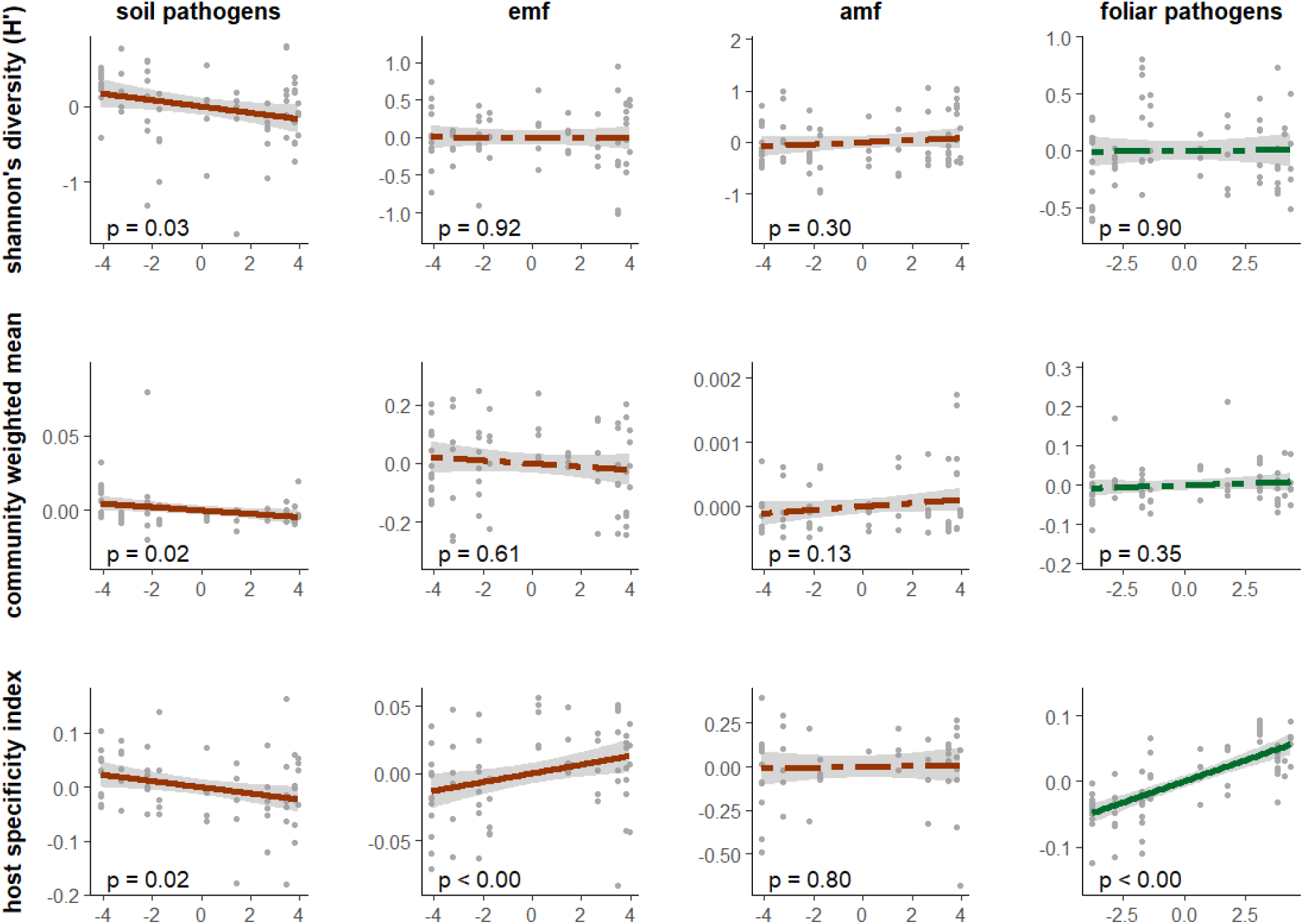
Soil pathogens richness, relative abundance and host specificity decline with the climate and elevation gradient. Conversely, EMF host specificity increases with climate and elevation. Each figure represents a linear regression of either the Shannon’s diversity, the community weighted mean (CWM), or the host specificity associated with each fungal guild (EMF = ectomycorrhizae, AMF = arbuscular mycorrhizae). All x-axes represent the ‘climate’ variable, which is directly related to elevation and with smaller values associated with lower elevation sites and larger values associated with higher elevation sites. The y-axis represents the residual variation of the regression analyses testing all variables except for ‘climate’ (nutrients, tree host species, pH) against these guild attributes. Figures with significant relationships have solid lines (p < 0.05), while those with insignificant relationships have dashed line.

For EMF, host specificity positively correlated with climate PCA axis 1 (β = 0.005, p = 0.001), indicating an increase in host specificity for EMF taxa across the elevation gradient (Figure 3, Appendix S1: Table S4). However, our models did not support an association between climate and EMF alpha diversity or EMF CWM values (Figure 3, Appendix S1: Table S4). Instead, EMF diversity was associated with tree host species, and increasing soil pH (β = 0.22, p = 0.021) (Appendix S3: Figure S3). Contrastingly, EMF host specificity decreased with soil pH (β = –0.04, p < 0.000) (Appendix S3: Figure S3). These findings partially support our hypothesis by demonstrating mutualist host specificity to correlate with the climate gradient.

For AMF, we found no significant associations between climate PCA axis 1 and any of our three metrics for assessing guild abundance (Figure 3, Appendix S1: Table S5). Instead, AMF diversity was structured by tree host species, with greater diversity on TABR than on PSME or TSHE hosts (Appendix S3: Figure S3). AMF CWM values were also structured by tree host species, with a higher AMF CWM on PSME and TABR than on TSHE (Appendix S3: Figure S3). These findings do not support our hypothesis that mutualist accumulation is greater at high elevations relative to low elevations.

For foliar pathogens, host specificity positively correlated with climate PCA axis 1 (β = 0.02, p < 0.000), indicating an increase in host specificity across the elevation gradient (Figure 3, Appendix S1: Table S7). Neither foliar pathogen diversity nor foliar pathogen CWM values were significantly associated with climate. Foliar pathogen diversity (β = 0.30, p = 0.002) and CWM values (β = 0.032, p = 0.048) instead positively correlated with pH (Appendix S3: Figure S4). These findings do not support our hypothesis that foliar pathogens decline in host specificity and abundance with increasing elevation. However, we did find that the relative read abundance of *Nothophaeocryprocus gaeumanii,* a dominant foliar pathogen associated with PSME, marginally covaries with climate (β = –109.40, p = 0.072), with greater relative abundance associated with smaller climate values and thus at lower elevation sites (Figure 4).

**Figure 4.**
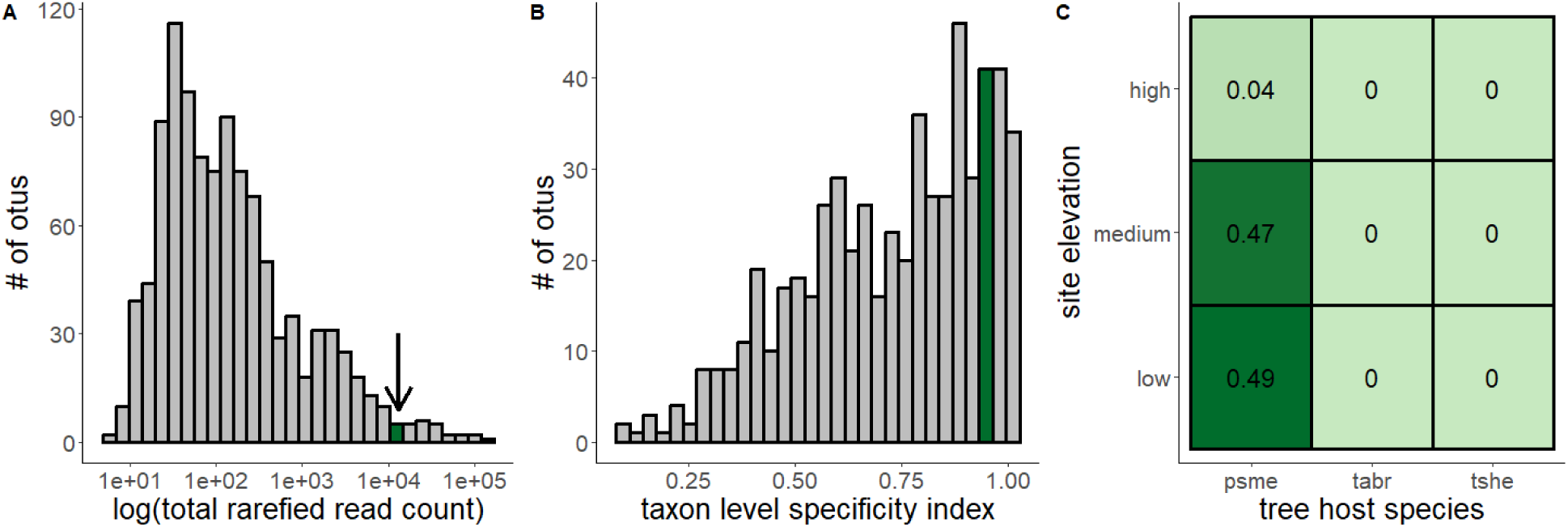
*Nothophaeocryptocus gaeumanii* is an abundant and host-specific foliar pathogen. Histogram of the number of total rarefied reads associated with each taxon in the dataset, with *N. gaeumanii* shown in green (A). Histogram of the specificity indices associated with each taxon in the data set, again, with *N. gaeumanii* shown in green (B). Heat map showing the greatest proportion of *N. gaeumanii* reads associated with PSME at low elevation sites (C).

## Discussion

### Host specificity of soil fungal functional groups correlates with elevation

Plant community diversity erodes as climates transition from warm and wet to dry and arid (Gentry 1988; Givnish 1999), and these patterns in plant species diversity across climate gradients are thought to be a result of variation in intraspecific competition. In relatively mesic environments, conspecific plants are hypothesized to accumulate shared enemies, which increase the fitness costs to growing near a conspecific plant, and result in more diverse plant communities (Janzen 1970, Connell 1971, Schemske et al 2009, LaManna 2022). Plant community diversity and plant-pathogenic microbial interactions have both independently been found to increase with precipitation (Gentry 1988; Givnish 1999; Curriero et al. 2001; Spear, Coley, and Kursar 2015). However, studies observing plant community diversity and microbial pathogens to covary with the same climate gradient are limited (Milici et al. 2020), and the role of mutualists in facilitating this diversity is even less explored (Zahra, Novotny, and Fayle 2021).

Here we examined fungal functional groups across a regional temperate elevation gradient known to negatively covary with CNDD strength (LaManna et al 2022). We found soil pathogen alpha diversity, relative abundance, and host specificity to decline with climate and elevation. These findings support our hypothesis that plants found at low elevations exhibit more host-specific pathogen accumulation relative to those found at high elevations. Moreover, by demonstrating soil pathogens negatively covary with the same climate gradient as does strength of CNDD and plant species diversity (LaManna et al 2022), our study represents a step in associating fungal pathogens with plant diversity in different environments.

The Janzen-Connell hypothesis (JCH) focuses on natural enemies mitigating plant community diversity patterns, and in the original text Connell (1971) postulates that forests without enemies would exist as single species groves with minimal intermixing (Connell 1971). A version of these single species groves theorized by Connell do exist naturally as monodominant stands in tropical forests (Connell and Lowman 1989), which otherwise harbor hyperdiverse plant communities (Gentry 1988). These monodominant stands have been hypothesized to exist because host-specific ectomycorrhizal mutualists may facilitate competitive exclusion of a single plant species (Connell and Lowman 1989; McGuire 2007), rather than as forest stands devoid of natural enemies. However, studies exploring the JCH, CNDD and microbes have mainly focused on soilborne pathogens (Bagchi et al. 2014; Bell, Freckleton, and Lewis 2006; Mangan et al. 2010), and evidence of mutualists contributing to patterns in CNDD remain limited (Zahra, Novotny, and Fayle 2021). We found EMF host specificity, but not alpha diversity or relative abundance, to increase across the climate gradient. These findings partially support our predictions and suggest that EMF may play a role in conspecific recruitment and thus the weakening of CNDD observed in the higher elevation forests. More broadly, we demonstrate that both soil pathogens and EMF communities exhibit variation in host specificity with climate. Pathogens cannot be implicated in intraspecific apparent competition, and likewise EMF cannot be implicated in intraspecific recruitment, without evidence of host specialization. By observing differences in host specialization in different environmental contexts, our results suggest that both guilds may independently contribute to CNDD strength. However, it has been proposed that extreme host specialization is not always a requirement to maintain plant species coexistence (Sedio and Ostling 2013) and multi-host pathogens may facilitate similar levels of plant diversity relative to pathogens that specialize on a single host (Benítez et al. 2013). In our study though, host specificity is considered a requirement, with the acknowledgement that generalist enemies may sometimes facilitate strong CNDD that can be misattributed to enemies that are highly host specific.

We found no relationship between AMF communities and our three metrics for evaluating guilds across the climate and elevation gradient. In our predictions, mutualists were grouped into one category, so the mutualist guilds (AMF and EMF) in our study were thus expected to exhibit similar trends in diversity, relative abundance and host specificity across the gradient. The category ‘mutualist’ is limiting given that AMF and EMF display distinct life histories and occupy different environmental niches and host species. AMF are widely considered to be generalist symbionts relative to EMF (Allen et al. 1995), and plant species associating with AMF symbionts are more sensitive to CNDD compared to those associating with EMF (Bennett et al. 2017; Pu et al. 2022). Further, AMF are more abundant in moist, mesic environments and EMF in cold, dry environments across global (Steidinger et al. 2019) and continental (Van Nuland et al. 2023) scales. Thus, it is perhaps unsurprising that we found that these two guilds are structured differently across the gradient. We only had one AMF-associated fungal host in our study (*Taxus brevifolia*), which appears to form mutualisms with both AMF and EMF. We found AMF community diversity and relative abundance to be driven by tree host, with *T. brevifolia* focal trees supporting a greater diversity relative to the other two focal tree species. These compositional differences attributed to tree host are strong, and may have prevented us from detecting a climate signal at the relatively small spatial scale examined in our study. Future studies that use a less targeted sampling approach and characterize soil communities representative of an entire site rather than associated with specific tree hosts could address the extent to which there is a trade-off in AMF and EMF fungi niches across a local climate gradient similar to those documented at larger scales (Steidinger et al. 2019; Van Nuland et al. 2023). Lastly, we were only able to classify 11 OTUs as AMF in the data set, a potential explanation for any lack of association with climate. This small sample size can likely be attributed to using non-AMF specific primers for metabarcoding (Lekberg et al. 2018), or a sampling scheme of collecting bulk soils rather than rhizosphere samples which selects for Ascomycota pathogens and saprobes over EMF and AMF species (Ren et al. 2021).

### Foliar fungal pathogens do not support our hypotheses through a community analysis, though a key foliar pathogen Nothophaeocryptocus gaeumanii does independently support our hypothesis

Although there are many examples of foliar microbial pathogens regulating plant populations in monoculture forest stands, agriculture, and nurseries (Grünwald, Goss, and Press 2008; Yarwood 1957), little is known about the role of foliar microbes in mitigating plant species diversity and distributions in natural ecosystems (Busby et al. 2022). Here we incorporate foliar fungi into our study design to investigate the extent to which aboveground microbes might contribute to CNDD strength. We focus on foliar pathogens, rather than both foliar pathogens and mutualists, in our guild level analyses because foliar mutualists are comparatively understudied in conifer systems (Sieber 2007). Foliar pathogen host specificity increased with the climate and elevation gradient, a result that is contrary to our hypothesis and opposite of what we found with soil pathogens. We also found no relationship between climate and either foliar pathogen diversity or relative abundance. However, a major challenge for interpreting these results is an incomplete understanding of foliar fungal function (Arnold 2007). The genetic and trait database infrastructure are not as well established for foliar communities relative to soil fungal communities, and the database limitations will influence downstream results and biological interpretations. For example, *Rhabdocline* is a genus that is dominant in the PSME foliar endophytic fungal microbiome (Gervers et al. 2022). In our data set, we identified two species in the genus *Rhabdocline*; *Rhabdocline weirii*, a pathogen known to be the causal agent of the disease Rhabdocline needle cast, and *Rhabdocline parkerii*, a putative mutualist (Carroll 1988). In both the FungalTraits and FUNGuild databases, fungi are sorted into guilds based on genus (Nguyen et al. 2016; Põlme et al. 2020), and the genus *Rhabdocline* is classified as a pathogen. Consequently, in our analyses, *Rhabdocline parkerii,* the putative mutualist, is instead classified as a pathogen. Although this limitation of the trait databases is not unique to foliar fungi, misclassifications will likely occur more frequently in fungal communities of which functional roles are not clearly defined or functional roles are more likely to vary within a genus. These are attributes of foliar endophytic communities, in which distinctions between mutualism and pathogen are often less clear relative to the belowground fungal functional guilds that we investigated in this analysis (Carroll 1988; Sieber 2007).

An alternative way to address our predictions with foliar fungi is to focus on specific taxa meeting the criteria put forth in the hypotheses, rather than analyzing the community as a whole. *Nothophaeocryptocus gaeumannii* (*N. gaeumannii*) is a fungal pathogen and causal agent for Swiss needle cast (SNC), a devastating disease for Pacific Northwestern forests and the timber industry (Hansen et al. 2000). *N. gaeumanii* is also known from previous studies to dominate the PSME foliar fungal community (Gervers et al. 2022). When focusing our analyses on *N. gaeumanii*, we found it to be a dominant and highly host specific taxa in the dataset with over 99% of its rarefied reads associated with PSME needles (Figure 4). We also found the relative abundance of *N. gaeumanii* to be negatively associated with the climate and elevation gradient (Figure 4). Thus consistent with our predictions, as *N. gaeumanii* is a dominant host-specific foliar pathogen that is enriched at lower elevation sites.

## Conclusion

Next-generation sequencing technology has enabled us to examine microbial communities across spatial scales that would be near impossible during the time that Janzen and Connell were conducting research. These tools reveal powerful, landscape level patterns in microbial community composition, which can have implications for intraspecific plant interactions and plant community diversity. However, pattern recognition is just the first step of the scientific process and, in our case, manipulative experiments are a necessary follow up to strengthen connections between the plant-associated microbiome, intraspecific plant interactions, and climate.

## Acknowledgements

Thank you to Bruce McCune for help with development of the specificity index. This work was supported by the US National Science Foundation under LTER8 DEB-2025755 (PEB), DEB 1542681 (FAJ), DEB 2310101 (PEB and FAJ), CAREER 2146552 (PEB), the US Department of Agriculture National Institute of Food and Agriculture under 2023-11569 (ASN and PEB), and CAREER 2022-67013-37437 (PEB). Any opinions, findings, and conclusions or recommendations expressed in this material are those of the authors and do not necessarily reflect the views of the National Science Foundation.

## Author Contributions

FAJ, FEA, PEB and ASN conceived the ideas tested in this study. ASN analyzed data and wrote the first draft of the MS. FAJ and FEA designed the study, collected and analyzed data. KAG assisted with data analysis. All authors contributed to the writing and editing of the MS.

