## Appendix 1 for "Host specificity of fungal functional groups covaries with elevation: implications for intraspecific plant interactions"

*Content of Appendix S1:* Tables of richness/diversity regression results and PERMANOVA results for the whole fungal communities; tables of regression results associated with climate and each fungal guild richness, diversity, and host specificity. The AICc model selection can be found in Appendix S2.
*Abbreviations for Appendix S1:* Df - degrees of freedom; SS - sum of squares, β - y intercept; SE - standard error; TSHE - *Tsuga heterophylla*; TABR - *Taxus brevifolia*

**Appendix S1: Table S1. Regression results display the relationship between climate and either richness or diversity for the soil and foliar fungal communities.**

| | β | SE | t-stat | P |
|---|---|---|---|---|
| **Soil Richness** | | | | |
| (Intercept) | 410.574 | 8.818 | 46.561 | 0.000 |
| climate | -9.346 | 2.836 | -3.296 | 0.002 |
| **Soil Diversity** | | | | |
| (Intercept) | 3.731 | 0.058 | 64.829 | 0.000 |
| climate | -0.027 | 0.019 | -1.476 | 0.145 |
| **Foliar Richness** | | | | |
| (Intercept) | 315.662 | 7.411 | 42.596 | 0.000 |
| climate | -7.490 | 2.400 | -3.121 | 0.003 |
| **Foliar Diversity** | | | | |
| (Intercept) | 3.450 | 0.044 | 79.049 | 0.000 |
| climate | 0.008 | 0.014 | 0.593 | 0.555 |

**Appendix S1: Table S2. PERMANOVA results display the amount of variation explained by climate and/or tree host species for soil and foliar communities.**

| | Df | SS | $R^2$ | F-stat | P |
|---|---|---|---|---|---|
| **Soil Permanova** | | | | | |
| climate | 1.000 | 1.774 | 0.075 | 5.344 | 0.001 |
| Residual | 66.000 | 21.912 | 0.925 | NA | NA |
| Total | 67.000 | 23.686 | 1.000 | NA | NA |
| **Foliar Permanova** | | | | | |
| tree host species | 2.000 | 2.615 | 0.170 | 6.827 | 0.001 |
| climate | 1.000 | 1.048 | 0.068 | 5.471 | 0.001 |
| Residual | 61.000 | 11.680 | 0.760 | NA | NA |
| Total | 64.000 | 15.368 | 1.000 | NA | NA |

**Appendix S1: Table S3. Regression results display the relationship between climate and soil pathogen community diversity, CWM, and host specificity.**

| | β | SE | t-stat | P |
|---|---|---|---|---|
| **Soil Pathogens** | | | | |
| *Diversity* | | | | |
| (Intercept) | 2.017 | 0.057 | 35.110 | 0.000 |
| climate | -0.067 | 0.018 | -3.631 | 0.001 |
| *CWM* | | | | |
| (Intercept) | 0.010 | 0.002 | 6.587 | 0.000 |
| climate | -0.002 | 0.001 | 3.147 | 0.002 |
| soil properties | 0.000 | 0.000 | 0.603 | 0.547 |
| *Host Specificity* | | | | |
| (Intercept) | 0.607 | 0.007 | 83.940 | 0.000 |
| climate | -0.008 | 0.002 | -3.493 | 0.001 |

**Appendix S1: Table S4. Regression results display the relationship between climate and EMF community diversity, CWM, and host specificity.**

| | | | | |
|---|---|---|---|---|
| **Soil EMF** | | | | |
| *Diversity* | | | | |
| (Intercept) | 1.056 | 0.509 | 2.077 | 0.042 |
| tree host (TABR) | 0.361 | 0.123 | 2.926 | 0.005 |
| tree host (TSHE) | 0.417 | 0.125 | 3.344 | 0.001 |
| soil pH | 0.222 | 0.093 | 2.373 | 0.021 |
| *CWM* | | | | |
| (Intercept) | 0.533 | 0.017 | 31.632 | 0.000 |
| soil properties | 0.000 | 0.000 | -0.954 | 0.343 |
| *Host Specificity* | | | | |
| (Intercept) | 0.798 | 0.044 | 18.243 | 0.000 |
| tree host (TABR) | 0.021 | 0.010 | 2.102 | 0.040 |
| tree host (TSHE) | 0.002 | 0.010 | 0.210 | 0.835 |
| soil pH | -0.035 | 0.008 | -4.385 | 0.000 |
| climate | 0.005 | 0.001 | 3.537 | 0.001 |

**Appendix S1: Table S5. Regression results display the relationship between climate and AM community diversity, CWM, and host specificity.**

| | β | SE | t-stat | P |
|---|---|---|---|---|
| **Soil AM** | | | | |
| *Diversity* | | | | |
| (Intercept) | -0.338 | 0.773 | 0.432 | 0.666 |
| tree host (TABR) | 0.440 | 0.165 | 2.625 | 0.009 |
| tree host (TSHE) | -0.167 | 0.167 | 0.981 | 0.327 |
| soil pH | 0.137 | 0.142 | 0.956 | 0.339 |
| climate | 0.017 | 0.023 | 0.723 | 0.470 |
| soil properties | 0.000 | 0.000 | 0.284 | 0.776 |
| *CWM* | | | | |
| (Intercept) | 0.000 | 0.001 | 0.278 | 0.781 |
| tree host (TABR) | 0.000 | 0.000 | 0.312 | 0.755 |
| tree host (TSHE) | 0.000 | 0.000 | 2.144 | 0.032 |
| soil pH | 0.000 | 0.000 | 0.790 | 0.430 |
| climate | 0.000 | 0.000 | 0.741 | 0.459 |
| *Host Specificity* | | | | |
| (Intercept) | 0.528 | 0.366 | 1.410 | 0.158 |
| tree host (TABR) | 0.063 | 0.087 | 0.705 | 0.481 |
| tree host (TSHE) | -0.093 | 0.095 | 0.959 | 0.337 |
| soil pH | 0.035 | 0.065 | 0.522 | 0.602 |
| soil properties | 0.000 | 0.000 | 0.420 | 0.674 |
| climate | 0.005 | 0.010 | 0.437 | 0.662 |

**Appendix S1: Table S6. Regression results display the relationship between climate and soil saprobe community diversity, CWM, and host specificity.**

| | β | SE | t-stat | P |
|---|---|---|---|---|
| **Soil Saprobes** | | | | |
| *Diversity* | | | | |
| (Intercept) | 3.852 | 0.739 | 5.209 | 0.000 |
| soil pH | -0.114 | 0.129 | -0.881 | 0.382 |
| *CWM* | | | | |
| (Intercept) | -0.117 | 0.115 | -1.019 | 0.312 |
| soil pH | 0.070 | 0.020 | 3.473 | 0.001 |
| *Host Specificity* | | | | |
| (Intercept) | 0.488 | 0.034 | 14.138 | 0.000 |

| | | | | |
|---|---|---|---|---|
| tree host (TABR) | 0.006 | 0.008 | 0.756 | 0.450 |
| tree host (TSHE) | 0.012 | 0.008 | 1.472 | 0.141 |
| soil pH | 0.021 | 0.006 | 3.326 | 0.001 |
| climate | 0.004 | 0.001 | 3.542 | 0.000 |
| soil properties | 0.000 | 0.000 | 0.432 | 0.666 |

**Appendix S1: Table S7. Regression results display the relationship between climate and foliar pathogen community diversity, CWM, and host specificity.**

| | β | SE | t-stat | P |
|---|---|---|---|---|
| **Foliar Pathogens** | | | | |
| *Diversity* | | | | |
| (Intercept) | 0.258 | 0.526 | 0.482 | 0.630 |
| tree host (TABR) | 0.108 | 0.124 | 0.861 | 0.389 |
| tree host (TSHE) | -0.094 | 0.120 | 0.773 | 0.439 |
| soil pH | 0.303 | 0.095 | 3.144 | 0.002 |
| foliar nutrients | 0.000 | 0.000 | 1.138 | 0.255 |
| *CWM* | | | | |
| (Intercept) | -0.050 | 0.085 | 0.574 | 0.566 |
| soil pH | 0.032 | 0.016 | 1.982 | 0.048 |
| tree host (TABR) | -0.022 | 0.020 | 1.080 | 0.280 |
| tree host (TSHE) | -0.037 | 0.025 | 1.473 | 0.141 |
| climate | 0.001 | 0.002 | 0.489 | 0.625 |
| *Host Specificity* | | | | |
| (Intercept) | 0.811 | 0.053 | 15.362 | 0.000 |
| tree host (TABR) | 0.057 | 0.012 | 4.746 | 0.000 |
| tree host (TSHE) | 0.024 | 0.012 | 2.065 | 0.043 |
| climate | 0.015 | 0.001 | 10.521 | 0.000 |
| soil pH | -0.022 | 0.010 | -2.244 | 0.029 |

**Appendix S1: Table S8. Regression results display the relationship between climate and foliar saprobe community diversity, CWM, and host specificity.**

| | β | SE | t-stat | P |
|---|---|---|---|---|
| **Foliar Pathogens** | | | | |
| *Diversity* | | | | |

| | | | | |
|---|---|---|---|---|
| (Intercept) | 1.195 | 0.675 | 1.769 | 0.082 |
| tree host (TABR) | 0.105 | 0.161 | 0.653 | 0.516 |
| tree host (TSHE) | -0.389 | 0.153 | -2.546 | 0.013 |
| soil pH | 0.191 | 0.124 | 1.540 | 0.129 |
| *CWM* | | | | |
| (Intercept) | 0.556 | 0.141 | 3.946 | 0.000 |
| tree host (TABR) | -0.090 | 0.033 | -2.708 | 0.009 |
| tree host (TSHE) | 0.009 | 0.034 | 0.271 | 0.787 |
| soil pH | -0.030 | 0.026 | -1.176 | 0.244 |
| foliar nutrients | 0.000 | 0.000 | 2.729 | 0.008 |
| *Host Specificity* | | | | |
| (Intercept) | 0.686 | 0.007 | 100.093 | 0.000 |
| tree host (TABR) | 0.042 | 0.009 | 4.615 | 0.000 |
| tree host (TSHE) | 0.004 | 0.009 | 0.399 | 0.691 |
| climate | -0.002 | 0.001 | -1.358 | 0.180 |
