## Appendix 2 for "Host specificity of fungal functional groups covaries with elevation: implications for intraspecific plant interactions"

*Content of Appendix S2:* AICc model selection associated with PERMANOVA models and regression analyses.

**Appendix S2: Table S1. AICc model selection associated with soil and foliar PERMANOVA analyses**

| | AICc | Delta AICc | Relative Likelihood | Model Weights |
|---|---|---|---|---|
| **Soil PERMANOVA** | | | | |
| Host Species + Climate + Soil pH + Soil Properties | -71.208 | 0.512 | 0.774 | 0.243 |
| Host Species + Climate + Soil pH | -72.083 | 0.364 | 0.834 | 0.262 |
| Host Species + Climate | -71.719 | 1.106 | 0.575 | 0.181 |
| Climate | -72.826 | 0.000 | 1.000 | 0.314 |
| **Foliar PERMANOVA** | | | | |
| Host Species + Climate + Soil pH + Foliar Properties | -102.510 | 0.395 | 0.821 | 0.283 |
| Host Species + Climate + Soil pH | -102.657 | 0.249 | 0.883 | 0.305 |
| Host Species + Climate | -102.906 | 0.000 | 1.000 | 0.345 |
| Host Species | -99.596 | 3.309 | 0.191 | 0.066 |

**Appendix S2: Table S2. AICc model selection associated with the regression model exploring the relationship between climate and soil pathogen community diversity, CWM, and host specificity.**

| | AICc | Delta AICc | Log-Likelihood | Relative Likelihood | Model Weight |
|---|---|---|---|---|---|
| **Soil Pathogens** | | | | | |
| *Diversity* | | | | | |
| Host Species + Climate + Soil pH + Soil Properties | 102.354 | 6.574 | -43.244 | 0.037 | 0.026 |
| Host Species + Climate + Soil Properties | 101.343 | 5.563 | -43.983 | 0.062 | 0.043 |
| Climate + Soil Properties | 97.911 | 2.132 | -44.638 | 0.344 | 0.239 |
| Climate | 95.780 | 0.000 | -44.702 | 1.000 | 0.693 |
| *CWM* | | | | | |
| Host Species + Climate + Soil pH + Soil Properties | -392.025 | 7.479 | 203.946 | 0.024 | 0.013 |
| Host Species + Climate + Soil Properties | -394.508 | 4.996 | 203.942 | 0.082 | 0.044 |
| Climate + Soil Properties | -398.949 | 0.555 | 203.792 | 0.758 | 0.407 |
| Climate | -399.504 | 0.000 | 202.940 | 1.000 | 0.537 |

| | | | | | |
|---|---|---|---|---|---|
| *Host Specificity* | | | | | |
| Host Species + Climate + Soil pH + Soil Properties | -203.126 | 6.220 | 109.496 | 0.045 | 0.036 |
| Host Species + Climate + Soil Properties | -204.001 | 5.344 | 108.689 | 0.069 | 0.056 |
| Climate + Soil Properties | -204.946 | 4.399 | 107.957 | 0.111 | 0.091 |
| Climate | -209.346 | 0.000 | 107.860 | 1.000 | 0.817 |

**Appendix S2: Table S3. AICc model selection associated with the regression model exploring the relationship between climate and soil EMF community diversity, CWM, and host specificity.**

| | AICc | Delta AICc | Log-Likelihood | Relative Likelihood | Model Weight |
|---|---|---|---|---|---|
| **Soil EMF** | | | | | |
| *Diversity* | | | | | |
| Host Species + Climate + Soil pH + Soil Properties | 76.489 | 4.779 | -30.311 | 0.092 | 0.064 |
| Host Species + Soil pH + Soil Properties | 74.014 | 2.303 | -30.318 | 0.316 | 0.221 |
| Host Species + Soil pH | 71.710 | 0.000 | -30.371 | 1.000 | 0.700 |
| pH | 79.540 | 7.830 | -36.583 | 0.020 | 0.014 |
| *CWM* | | | | | |
| Host Species + Climate + Soil pH + Soil Properties | -64.939 | 6.132 | 40.403 | 0.047 | 0.036 |
| Host Species + Soil pH + Soil Properties | -65.989 | 5.082 | 39.683 | 0.079 | 0.060 |
| Host Species + Soil Properties | -67.664 | 3.407 | 39.316 | 0.182 | 0.139 |
| Soil Properties | -71.071 | 0.000 | 38.723 | 1.000 | 0.765 |
| *Host Specificity* | | | | | |
| Host Species + Climate + Soil pH + Soil Properties | -312.886 | 5.991 | 163.132 | 0.050 | 0.047 |
| Host Species + Soil pH + Climate | -318.877 | 0.000 | 167.372 | 1.000 | 0.990 |
| Host Species + Climate | -309.350 | 9.527 | 160.159 | 0.009 | 0.008 |
| Climate | -305.940 | 12.936 | 156.158 | 0.002 | 0.001 |

**Appendix S2: Table S4. AICc model selection associated with the regression model exploring the relationship between climate and soil AM community diversity, CWM, and host specificity.**

| | AICc | Delta AICc | Log-Likeliho | Relative | Model |
|---|---|---|---|---|---|

| | | | od | Likelihood | Weight |
|---|---|---|---|---|---|
| **Soil AM** | | | | | |
| *Diversity* | | | | | |
| Host Species + Climate + Soil pH + Soil Properties | 111.525 | 1.486 | -47.829 | 0.476 | 0.158 |
| Host Species + Soil pH + Climate | 110.039 | 0.000 | -48.331 | 1.000 | 0.332 |
| Host Species + Soil pH | 110.997 | 0.958 | -50.015 | 0.619 | 0.206 |
| Host Species | 110.215 | 0.177 | -50.790 | 0.915 | 0.304 |
| *CWM* | | | | | |
| Host Species + Climate + Soil pH + Soil Properties | -836.390 | 2.352 | 426.128 | 0.308 | 0.121 |
| Host Species + Soil pH + Climate | -838.742 | 0.000 | 426.060 | 1.000 | 0.392 |
| Host Species + Soil pH | -836.942 | 1.800 | 423.955 | 0.407 | 0.159 |
| Host Species | -838.380 | 0.362 | 423.507 | 0.834 | 0.327 |
| *Host Specificity* | | | | | |
| Host Species + Climate + Soil pH + Soil Properties | 3.124 | 1.309 | 6.838 | 0.520 | 0.362 |
| Host Species + Soil pH + Climate | 3.162 | 1.347 | 5.443 | 0.510 | 0.355 |
| Host Species + Soil pH | 3.614 | 1.798 | 3.907 | 0.407 | 0.283 |
| Host Species | 1.816 | 0.000 | 3.557 | 1.000 | 0.410 |

**Appendix S2: Table S5. AICc model selection associated with the regression model exploring the relationship between climate and soil saprobe community diversity, CWM, and host specificity.**

| | AICc | Delta AICc | Log-Likelihood | Relative Likelihood | Model Weight |
|---|---|---|---|---|---|
| **Soil Saprobes** | | | | | |
| *Diversity* | | | | | |
| Host Species + Climate + Soil pH + Soil Properties | 130.546 | 4.693 | -57.339 | 0.096 | 0.066 |
| Host Species + Soil pH + Climate | 128.564 | 2.712 | -57.593 | 0.258 | 0.177 |
| Host Species + Soil pH | 130.367 | 4.515 | -59.700 | 0.105 | 0.072 |
| Soil pH | 125.852 | 0.000 | -59.739 | 1.000 | 0.686 |
| *CWM* | | | | | |
| Host Species + Climate + Soil pH + Soil Properties | -118.567 | 8.377 | 67.217 | 0.015 | 0.013 |
| Host Species + Soil pH + Climate | -120.948 | 5.996 | 67.163 | 0.050 | 0.042 |

| | | | | | |
|---|---|---|---|---|---|
| Host Species + Soil pH | -122.649 | 4.295 | 66.809 | 0.117 | 0.099 |
| Soil pH | -126.944 | 0.000 | 66.660 | 1.000 | 0.846 |
| *Host Specificity* | | | | | |
| Host Species + Climate + Soil pH + Soil Properties | -301.433 | 1.443 | 158.650 | 0.486 | 0.324 |
| Host Species + Soil pH + Climate | -302.876 | 0.000 | 158.127 | 1.000 | 0.666 |
| Host Species + Climate | -294.055 | 8.821 | 152.512 | 0.012 | 0.008 |
| Climate | -291.532 | 11.344 | 148.954 | 0.003 | 0.002 |

**Appendix S2: Table S6. AICc model selection associated with the regression model exploring the relationship between climate and foliar pathogen community diversity, CWM, and host specificity.**

| | AICc | Delta AICc | Log-Likelihood | Relative Likelihood | Model Weight |
|---|---|---|---|---|---|
| **Foliar Pathogens** | | | | | |
| *Diversity* | | | | | |
| Host Species + Climate + Soil pH + Foliar Nutrients | 56.044 | 2.496 | -20.039 | 0.287 | 0.147 |
| Host Species + Soil pH + Foliar Nutrients | 53.548 | 0.000 | -20.050 | 1.000 | 0.512 |
| Soil pH + Foliar Nutrients | 56.189 | 2.642 | -23.761 | 0.267 | 0.137 |
| Soil pH | 55.388 | 1.840 | -24.497 | 0.398 | 0.204 |
| *CWM* | | | | | |
| Host Species + Climate + Soil pH + Foliar Nutrients | -187.272 | 2.310 | 101.618 | 0.315 | 0.130 |
| Host Species + Soil pH + Climate | -188.807 | 0.775 | 101.128 | 0.679 | 0.280 |
| Host Species + Soil pH | -189.582 | 0.000 | 100.299 | 1.000 | 0.413 |
| Soil pH | -187.887 | 1.695 | 97.140 | 0.429 | 0.177 |
| *Host Specificity* | | | | | |
| Host Species + Climate + Soil pH + Foliar Nutrients | -245.282 | 2.510 | 130.623 | 0.285 | 0.186 |
| Host Species + Soil pH + Climate | -247.791 | 0.000 | 130.620 | 1.000 | 0.653 |
| Host Species + Climate | -244.984 | 2.807 | 128.000 | 0.246 | 0.160 |
| Climate | -232.427 | 15.365 | 119.410 | 0.000 | 0.000 |

**Appendix S2: Table S7. AICc model selection associated with the regression model exploring the relationship between climate and foliar saprobe community diversity, CWM, and host specificity.**

| | AICc | Delta AICc | Log-Likelihood | Relative Likelihood | Model Weight |
|---|---|---|---|---|---|
| **Foliar Saprobes** | | | | | |
| *Diversity* | | | | | |
| Host Species + Climate + Soil pH + Foliar Nutrients | 92.862 | 4.610 | -38.449 | 0.100 | 0.070 |
| Host Species + Soil pH + Foliar Nutrients | 90.565 | 2.314 | -38.559 | 0.314 | 0.221 |
| Host Species + Soil pH | 88.251 | 0.000 | -38.617 | 1.000 | 0.704 |
| Soil pH | 98.518 | 10.267 | -46.062 | 0.006 | 0.004 |
| *CWM* | | | | | |
| Host Species + Climate + Soil pH + Foliar Nutrients | -113.016 | 2.099 | 64.490 | 0.350 | 0.256 |
| Host Species + Soil pH + Foliar Nutrients | -115.115 | 0.000 | 64.281 | 1.000 | 0.732 |
| Soil pH + Foliar Nutrients | -106.344 | 8.770 | 57.506 | 0.012 | 0.009 |
| Foliar Nutrients | -104.284 | 10.831 | 55.339 | 0.004 | 0.003 |
| *Host Specificity* | | | | | |
| Host Species + Climate + Soil pH + Foliar Nutrients | -267.794 | 4.55928 | 141.8797 | 0.102 | 0.072 |
| Host Species + Soil pH + Climate | -270.015 | 2.33825 | 141.7319 | 0.311 | 0.220 |
| Host Species + Climate | -272.354 | 0 | 141.6853 | 1.000 | 0.708 |
| Climate | -251.521 | 20.83311 | 128.957 | 0.000 | 0.000 |
