## Appendix 3 for "Host specificity of fungal functional groups covaries with elevation: implications for intraspecific plant interactions"

*Content of Appendix S3:* Figures associated with additional analyses between fungal guilds and environmental variables

**Appendix S3: Figure S1. Rarefaction curves show that fungal species richness is correlated with elevation, with greater richness at lower elevation sites for both soil fungal communities (A) and needle fungal communities (B).** Each curve represents the species accumulation for one sample in the dataset, with the x-axis representing the read depth of each sample, and the y-axis representing the number of OTUs in each sample. The color gradient signifies the elevation of the tree host species associated with each sample, and the gray vertical line signifies the minimum read depth used in both data sets (25,303) .

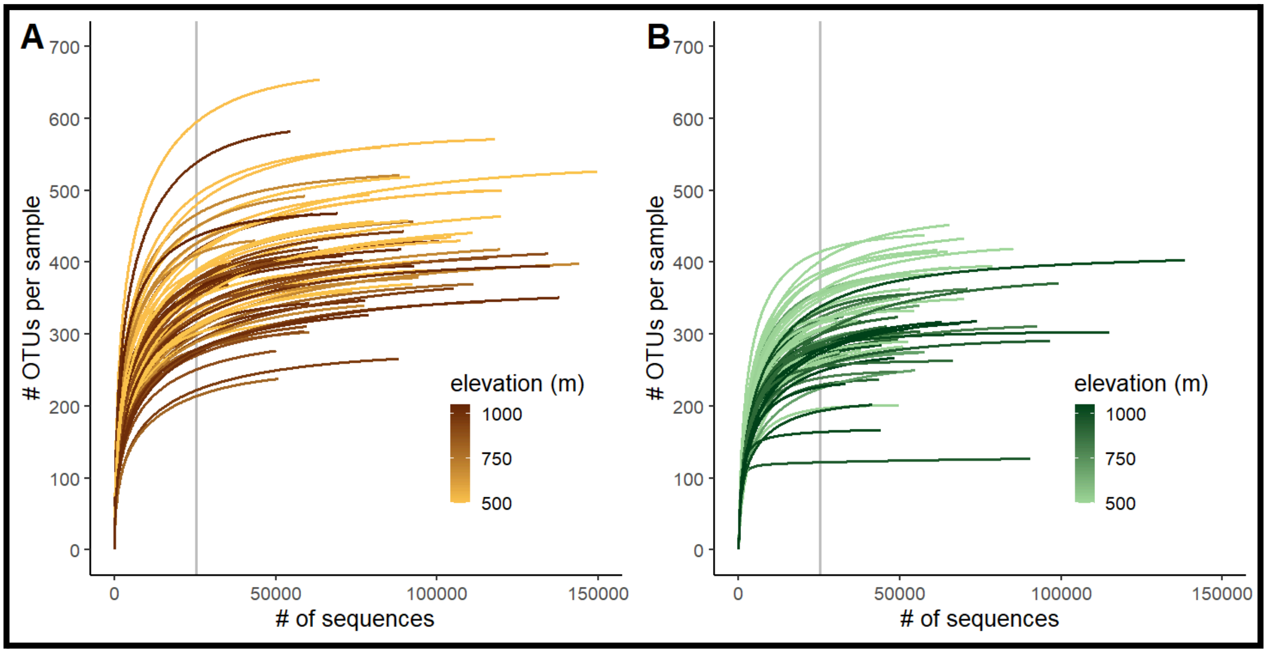

**Appendix S3: Figure S2. Principal Component Analysis of the climate (A), soil properties (B) and foliar nutrients (C) show the relative contributions of each attribute to the first dimension.** PC 1 was used for PERMANOVA and regression analyses. Fig. S2D demonstrates that the climate variable used in analyses is correlated with elevation.

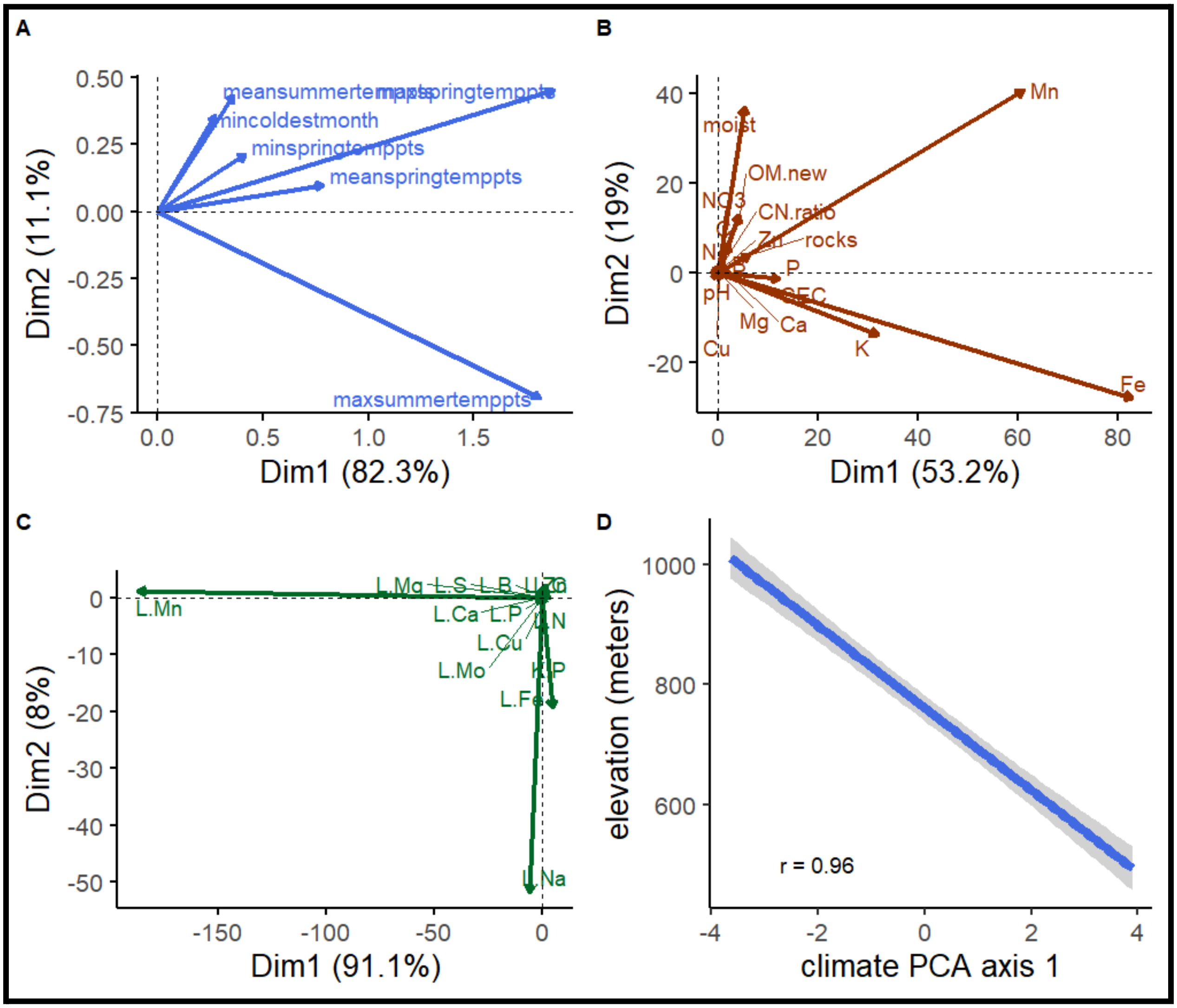

**Appendix S3: Figure S3. Linear regression analyses and ANOVA tests of diversity (A,B,D), host specificity (C), and CWMs (E,F) show how different soil fungal guilds are associated with abiotic and biotic attributes of the data set.** Only significant associations are shown here ($p < 0.1$), amf = arbuscular mycorrhizae, emf = ectomycorrhizae.

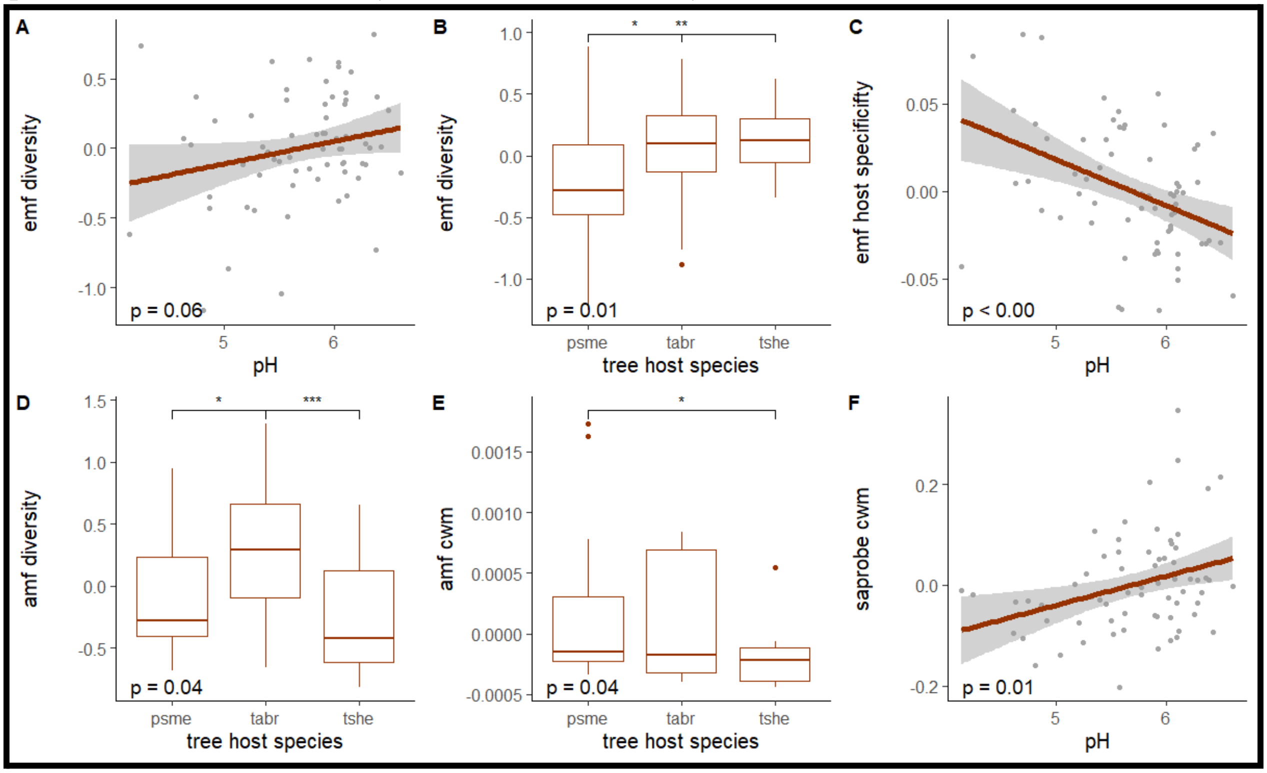

**Appendix S3: Figure S4. Linear regression analyses and ANOVA tests of diversity (A,D), CWMs (B,E), and host specificity (C,G) show how different foliar fungal guilds are associated with abiotic and biotic attributes of the data set.** Only significant associations are shown here ($p < 0.1$).

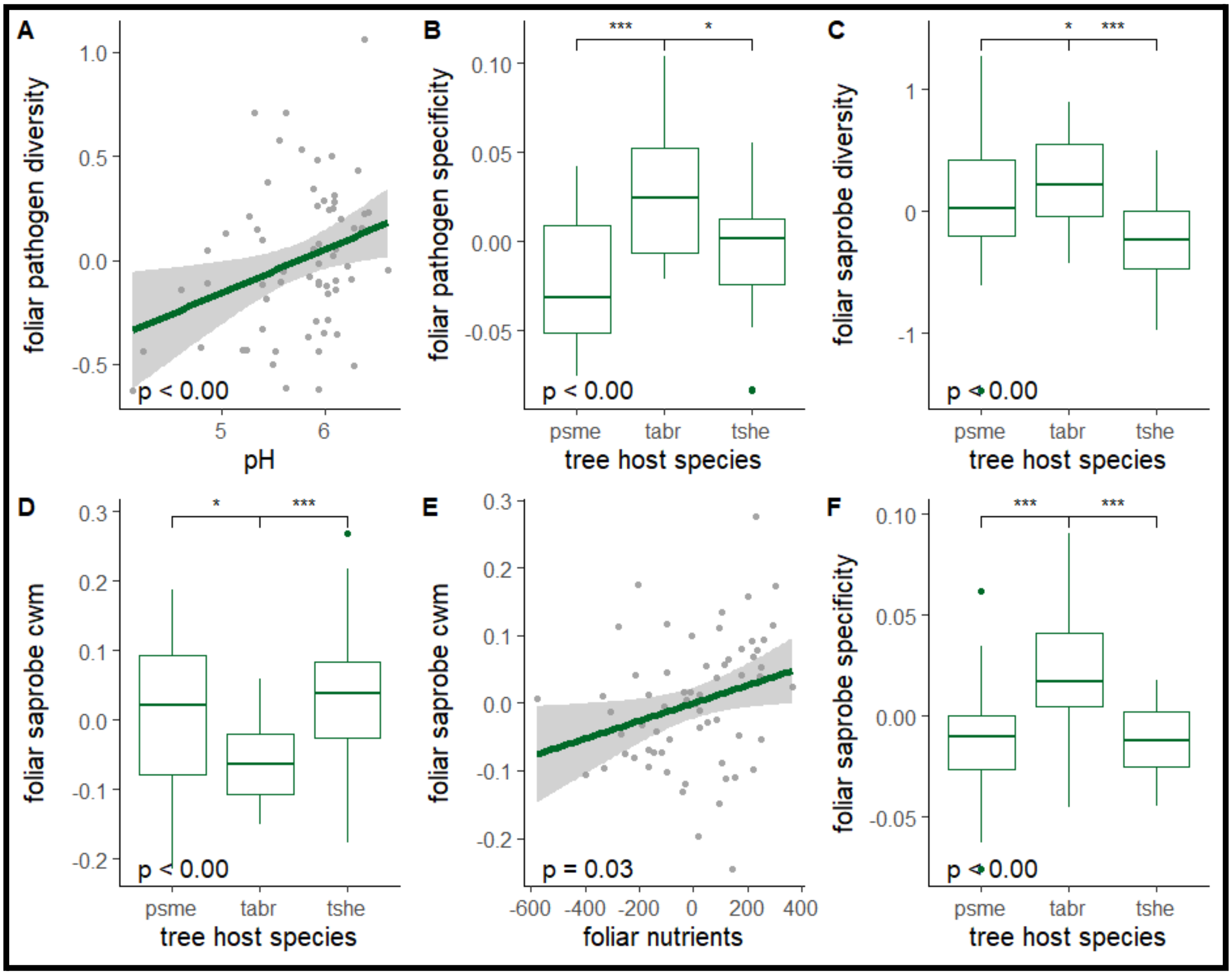
