## Appendix 4 for "Host specificity of fungal functional groups covaries with elevation: implications for intraspecific plant interactions"

*Content of Appendix S4:* detailed guide for specificity index calculations

Here we define host specificity as the degree to which a microbial species is associated with a certain host, or rather, is more widely associated with many host species (Christian, Whitaker, and Clay 2015). With this index, our goal was to determine a value between 0-1 that describes the extent to which a host-associated fungal community is host specific. To address this goal, we first ascribe each OTU a specificity value by using the results of the permutation test associated with the Indicator Species Analysis. We then treat these values as 'traits' associated with each taxa, and we use the Community Weighted Mean analysis to translate these species level values to specificity indices associated with each host in the dataset. To allow taxa to behave differently in different environments, we split the data into two and analyzed the data separately: high elevation focal trees and low elevation focal trees. Below we describe our calculations using example values from our low elevation foliar fungal community dataset.

1. We first performed the Indicator Species Analysis with the dataset.
   a. We split up the sample units into groups. In our study, each group represented a focal tree host. There are three groups total (PSME, TSHE, TABR).
   b. We then calculated the Indicator Value (IV) for each taxa associated with each group. The IV is found by multiplying a taxon's relative occurrence in each group [A] x its fidelity to each group [B] (i.e. given all the individuals in a group, this is the number of individuals that the taxon is actually found on), then dividing that number by 100 (Dufrêne and Legendre 1997). The resulting value was between 0 and 100, with taxa that are more likely to be an indicator species exhibiting values closer to 100. All analyses associated with the ISA were calculated using PC-Ord (McCune, 1986). See tables 1-3 for example calculations.

*Table 1. Calculations of the [A] relative occurrence of each taxon associated with each tree host.*

| otu id | relative occurrence PSME [A] | relative occurrence TSHE [A] | relative occurrence TABR [A] |
|---|---|---|---|
| **otu 1** | 0 | 0 | 100 |
| **otu 2** | 38 | 0 | 62 |
| **otu 3** | 1 | 92 | 7 |
| **otu 4** | 46 | 26 | 28 |
| **otu 5** | 20 | 21 | 59 |

*Table 2. Calculations of the [B] fidelity of each taxon associated with each tree host.*

| otu id | fidelity PSME [B] | fidelity TSHE [B] | fidelity TABR [B] |
|---|---|---|---|
| **otu 1** | 0 | 0 | 42 |
| **otu 2** | 10 | 0 | 8 |
| **otu 3** | 10 | 42 | 33 |
| **otu 4** | 30 | 25 | 8 |

| | | | |
|---|---|---|---|
| **otu 5** | 80 | 75 | 92 |

*Table 3. Calculation of Indicator Values [A] x [B]/100 of each taxon associated with each host.*

| otu id | Indicator Values (IVs) PSME [A] x [B] | Indicator Values (IVs) TSHE [A] x [B] | Indicator Values (IVs) TABR [A] x [B] |
|---|---|---|---|
| **otu 1** | 0 | 0 | 42 |
| **otu 2** | 4 | 0 | 5 |
| **otu 3** | 0 | 39 | 2 |
| **otu 4** | 14 | 7 | 2 |
| **otu 5** | 16 | 16 | 54 |

c. We then found the group with the highest IV, or $IV_{max}$. We used a Monte Carlo randomization significance test to determine how frequently this $IV_{max}$ will be higher than one generated by chance. This was accomplished by first randomizing the tree host sample units, calculating the $IV_{max}$ with the random sample unit assignments, then repeating this process iteratively. The resulting p-value represented the proportion of times that the randomized $IV_{max}$ was equal to or greater than the original $IV_{max}$ from the dataset (McCune and Grace 2002) (Table 4). A high $IV_{max}$ in combination with a significant result from this randomization test were attributes of a good indicator species.

*Table 4. Results of the randomization test.*

| otu id | IV max | standard deviation | p-value |
|---|---|---|---|
| **otu 1** | 42 | 7.23 | 0.005 |
| **otu 2** | 5 | 5.54 | 1.000 |
| **otu 3** | 39 | 10.20 | 0.163 |
| **otu 4** | 14 | 7.42 | 0.624 |
| **otu 5** | 54 | 5.56 | 0.007 |

2. We cannot not use Indicator Values (IVs) or the ISA directly for our specificity analysis because taxa that are highly host specific (i.e. high [A]) but in low abundance (i.e. low [B]) will have a low $IV_{max}$. Conversely, taxa that are very common (i.e. high [B]) but are not host specific (i.e. low [A]) will have a high $IV_{max}$ relative to their degree of host specificity. Thus, IVs are not a good representation of host specificity. We determined that the result of the randomization test of the $IV_{max}$ is a robust numerical representation of host specificity. We found that with each randomized distributions of $IV_{max}$, specialist taxa will have observed $IV_{maxes}$ on the extreme end of their associated distributions, whereas more generalist taxa will have $IV_{maxes}$ in the middle of their associated distributions (Figure 1). Attributes of taxa with insignificant results from the randomization test are common taxa that are generalists (these would have a high $IV_{max}$), rare taxa that are generalists (these would have a low $IV_{max}$), and rare taxa that are found in too few sites for statistical analyses (See Figure 2 below). Table 5 provides more information on how

taxa behavior influences both $IV_{max}$ values and the results of the randomization test (i.e. p-value of ISA).

**Figure 1. Conceptual figure of where a generalist IVmax (A) falls in the distribution of randomized IVmaxes vs a specialist IVmax (B).**

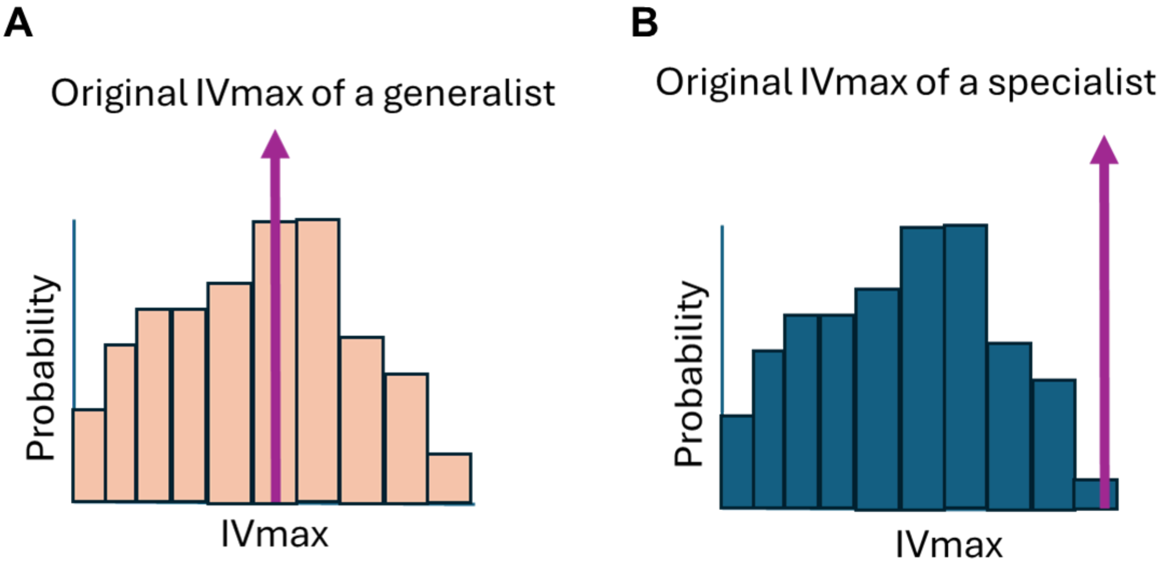


*Table 5. $IV_{max}$ values and p-values associated with taxa that range from generalist - specialist and rare - specific.*

| taxa behavior | relative abundance on dominant host [A] | fidelity on dominant host [B] | $IV_{max}$ | p-value |
|---|---|---|---|---|
| high abundance/host specific | high | high | high | low |
| high abundance/generalist | low | high | mid | high |
| low abundance/host specific | high | low | mid | low |
| low abundance/generalist | low | low | low | high |

3. In order to remove taxa that are too rare to exhibit a significant result from the randomization test, we made a figure to visualize the relationship between the p-values and the number of sites that each taxa is found in (Figure 2). We identified a threshold in which a taxa was in too few sites to have the potential of a significant p-value. We used this threshold (4 sites) as the cutoff point for rare taxa. All taxa found in less than 4 sites were excluded from downstream analyses.

**Figure 2. Scatter plot with each point representing an OTU in the dataset, with the x-axis indicating the number of sites that the OTU was found in and the y-axis the p-value from the randomization test.** The black horizontal line represents the significant p-value of 0.05 and the black vertical line represents our threshold of which to remove rare taxa, those found in less than 4 sites.

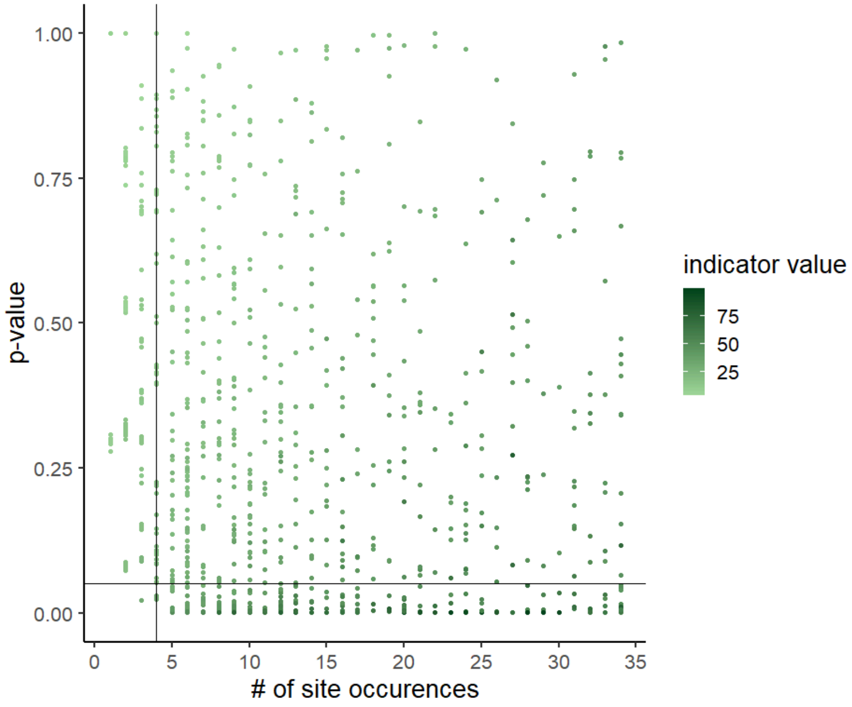


4. We subtracted the results of the randomization test (ie each p-value) from 1 to get a specificity value associated with each taxon in the data set. Low values indicate a generalist taxon and high values indicate a specialist taxon (Table 6).

*Table 6. Conversion of p-values into a specificity value associated with each taxon in the dataset.*

| otu id | p-value | 1-p |
|---|---|---|
| **otu 1** | 0.005 | 0.995 |
| **otu 2** | 1.000 | 0.000 |
| **otu 3** | 0.163 | 0.837 |
| **otu 4** | 0.624 | 0.376 |
| **otu 5** | 0.007 | 0.993 |

5. We then used the taxon level specificity values to calculate a community weighted mean (CWM) analysis. This generated specificity indices associated with each tree host in the data set, which represents the degree to which the community associated with it has specialist taxa. Higher values correspond to communities with more specialist taxa, and lower values correspond to communities with more generalist taxa.

6. We repeated steps 1-5 with both subsets of data, then merged the CWM values from the different subsets into one dataset.

7. Finally, we regressed these specificity indices against the independent variables (in this case climate, tree host species, foliar nutrients, soil pH) to determine if there is a relationship between site level specificity and climate.

<u>**References**</u>

Christian, Natalie, Briana Kathleen Whitaker, and Keith Clay. 2015. “Microbiomes: Unifying Animal and Plant Systems through the Lens of Community Ecology Theory.” *Frontiers in Microbiology* 6 (September). https://doi.org/10.3389/fmicb.2015.00869.

Dufrêne, Marc, and Pierre Legendre. 1997. “Species Assemblages and Indicator Species:The Need for a Flexible Asymmetrical Approach.” *Ecological Monographs* 67 (3): 345–66. https://doi.org/10.1890/0012-9615(1997)067[0345:SAAIST]2.0.CO;2.

McCune, B., and J. B. Grace. 2002. *Analysis of Ecological Communities*. https://www.waterbouwkundiglaboratorium.be/nl/publicaties/documentatiecentrum/catalogus.

McCune, Bruce. (1986). Pc-Ord: An Integrated System for Multivariate Analysis of Ecological Data. *Abstracta Botanica* 10, no. 2 : 221–25.
